# Activation of the mevalonate pathway in response to anti-cancer treatments drives glioblastoma recurrences through activation of Rac-1

**DOI:** 10.1101/2023.07.23.550205

**Authors:** Ling He, Angeliki Ioannidis, Evelyn Arambula, Carter J. Hoffman, Purva Joshi, Anoushka Kathiravan, Julian Whitelegge, Linda M. Liau, Harley I. Kornblum, Frank Pajonk

## Abstract

Glioblastoma is the deadliest adult brain cancer. Under the current standard of care almost all patients succumb to the disease and novel treatments are urgently needed. Dopamine receptor antagonists have been shown to target cancer cell plasticity in GBM and repurposing these FDA-approved drugs in combination with radiation improves the efficacy of radiotherapy in glioma models. In cells surviving this combination treatment the mevalonate pathway is upregulated at the transcriptional and functional level.

Here we report that glioblastoma treatments that converge in the immediate early response to radiation through activation of the MAPK cascade universally upregulate the mevalonate pathway and increase stemness of GBM cells through activation of the Rho-GTPase Rac-1. Activation of the mevalonate pathway and Rac-1 is inhibited by statins, which leads to improved survival in mouse models of glioblastoma when combined with radiation and drugs that target the glioma stem cell pool and plasticity of glioma cells.

## Introduction

Glioblastoma (GBM) is the deadliest adult brain cancer. The standard-of-care, surgery followed by radiation therapy and temozolomide (TMZ) almost always fails and the tumors progress or recur, thus leading to unacceptable low 5-year survival rates. Despite iterations in radiotherapy protocols and techniques, introduction of targeted therapies and biologics, the survival rates have not changed in almost two decades and strategies against GBM have hit a critical barrier. Understanding mechanisms underlying this treatment resistance holds the key to develop novel approaches against GBM.

GBMs are thought to be governed by a hierarchical organization of tumor cells with a small number of glioma stem cells (GSCs) at the top of this hierarchy, giving rise to the bulk of more differentiated glioma cells. GSCs have been reported to resist many chemotherapeutic agents [1] and are relatively resistant to ionizing radiation [2]. Consequently, a portion of the GSC population survives treatment and can repopulate the tumor. This led to efforts specifically targeting the GSC population of GBMs but so far this has not successfully translated into the clinic.

Recently, we reported an additional novel aspect of GBM biology in response to radiation. Non-stem glioma cells surviving irradiation converted into glioma-initiating cells in a radiation dose-dependent manner. Compounds that counteract this phenomenon improved median survival in patient-derived orthotopic xenograft (PDOX) and syngeneic models of GBM [3–5]. Yet, the efficacy was not 100% and bulk RNA-sequencing revealed that surviving cells upregulated gene expression of the mevalonate pathway, followed by upregulation of its enzymatic function *in vitro*. This hinted at a novel metabolic vulnerability that could be exploited by the addition of a statin, but it was unclear if the same effects could be observed *in vivo* and how the mevalonate pathway affects stemness of glioma cells. In the present study we report that combination treatments that synergize with radiation in activating the immediate early response through the mitogen-activated protein kinase (MAPK) cascade upregulate the mevalonate pathway *in vivo*, thereby triggering repopulation of gliomas through prenylation of the Rho GTPase Rac-1.

## Results

### Upregulation of the mevalonate pathway in response to anti-cancer treatments in vivo is restricted to glioma cells

We previously reported that combinations of radiation and dopamine receptor antagonists upregulated genes in the mevalonate pathway and increased cholesterol biosynthesis in GBM cells *in vitro* [3, 4]. Genes in this pathway are under control of sterol regulatory element-binding protein 2 (SREBP-2 also called SREBF-2), a transcription factor under control of the MAPK cascade though AP-1, a pathway well known to be activated during the immediate-early response to radiation [6]. Its components JunB, JunD, and FosB, are all part of the first-order regulatory network of SREBP-2. JunB and JunD have weak transactivation domains and can be considered repressors in the presence of the strong transcriptional activator FosB. All three are regulated by GSK-3 [7], the kinase downstream of dopamine receptors, thus explaining synergistic effects of radiation and dopamine receptor antagonists on the mevalonate pathway.

TMZ, part of the current standard-of-care against GBM is known to activate the MAPK cascade in GBM [8]. Likewise, vincristine, a component of the PCV chemotherapy regimen against GBM, activates this pathway [9]. Western blots of patient-derived HK374 and HK217 glioma cells **(Supplementary Table 1)** confirmed that radiation, quetiapine (QTP; a dopamine receptor antagonist), TMZ and vincristine activated the MAPK cascade in our model system (**Fig. 1a-c, Suppl. Fig. 1a/b**). Next, we tested if radiation in combination with TMZ, vincristine, or QTP would also up-regulate the mevalonate pathway. As expected, combining radiation and TMZ upregulated gene expression of key enzymes in this pathway (**Fig. 1d/e**). Similar results were found when combining radiation and vincristine (**Fig. 1f/g**). We have previously reported the upregulation of mevalonate pathway in HK374 cells upon combination treatment of radiation plus QTP [3]. Here, similar results were also observed in HK217 cells **(Suppl. Fig. 1c)**. This indicated that combination therapies that converge on the immediate-early response to radiation [10] through activation of the MAPK cascade universally upregulate the mevalonate pathway, thereby creating a metabolic vulnerability in cells surviving the sublethal DNA damage caused by radiation.

**Figure 1.**
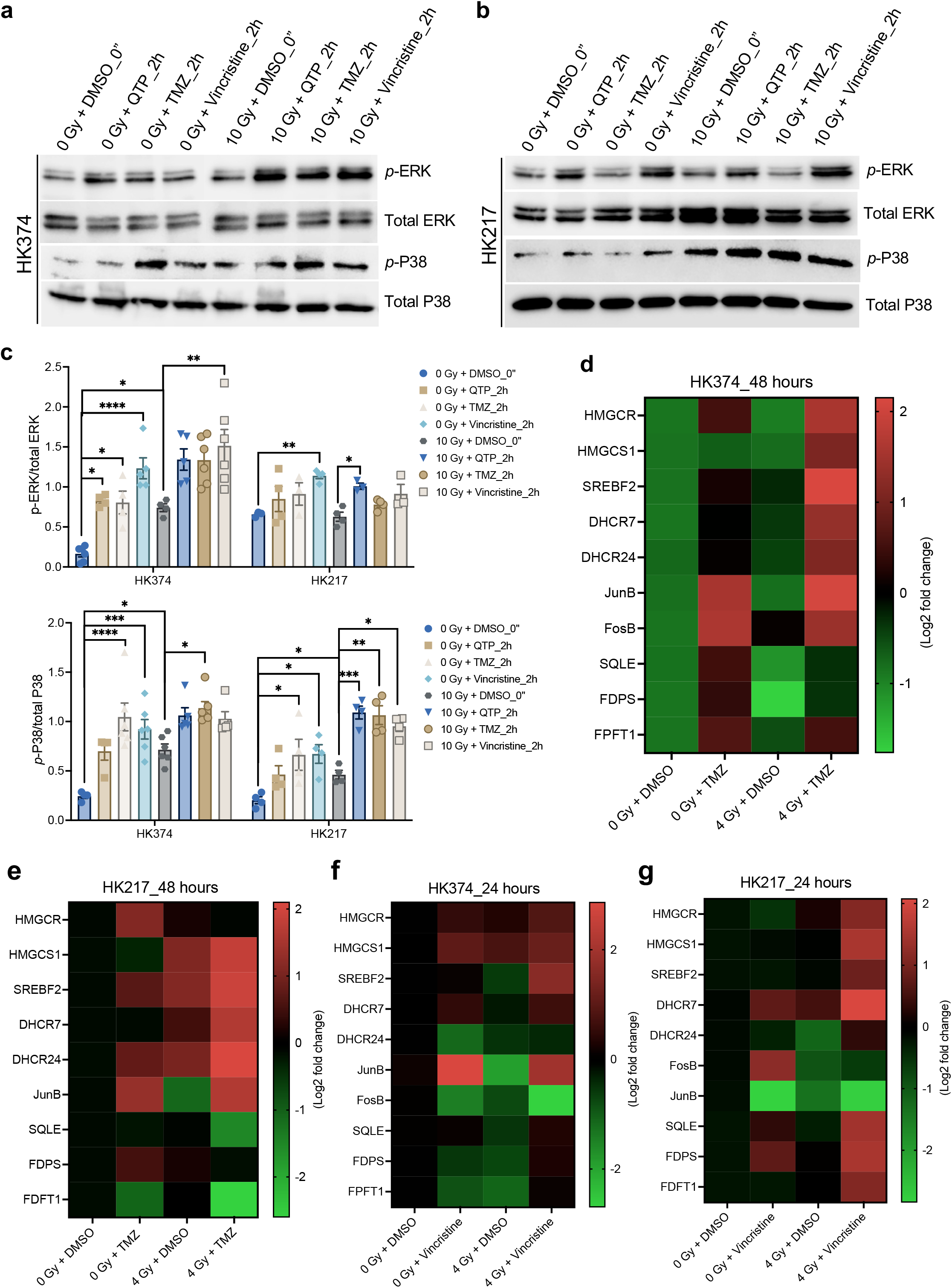
Combination therapies that converge on the immediate-early response to radiation through the MAPK cascade universally upregulate the mevalonate pathway. **(a/b)** Western blotting of *p*-ERK, total ERK, *p*-P38, and total P38 in patient-derived GBM HK374 and HK217 cell lines upon radiation (a single dose of 10 Gy) in combination of QTP (10 µM) or TMZ (1 mM) or Vincristine (250 nM) at 2 hours after treatment. (c) The densitometry measurements of *p*-ERK/total ERK and *p*-P38/total P38 using Image J. (d/e) Heatmap showing the results of quantitative RT-PCR for the cholesterol biosynthesis related genes in both HK374 and HK217 cells treated with radiation (a single dose of 4 Gy) in the presence of absence of TMZ (1 mM) for two consecutive days. (**f/g**) Heatmap showing the results of quantitative RT-PCR for the cholesterol biosynthesis related genes in both HK374 and HK217 cells treated with radiation (a single dose of 4 Gy) in the presence of absence of Vincristine (250 nM) for 24 hours. All experiments have been performed with at least 3 biological independent repeats. *p-*values were calculated using One-way ANOVA. * *p*-value < 0.05, ** *p*-value < 0.01, *** *p*-value < 0.001, **** *p*-value < 0.0001.

Our previous studies investigated the effects of radiation and dopamine receptor antagonists on the mevalonate pathway in patient derived GBM lines *in vitro* [3]. To test if this effect could also be observed in the presence of a tumor microenvironment, we next implanted HK374 glioma cells into the striatum of NSG mice. After 2 weeks of grafting and tumor growth, mice were treated with a single dose of 4 Gy and 2 consecutive doses of QTP **(Fig. 2a)**. Tumor tissue was harvested, mRNA extracted and subjected to qRT-PCR using human- and mouse-specific primers for genes in the mevalonate pathway. Gene expression of enzymes in the mevalonate pathway in tumors cells confirmed data obtained in our *in vitro* studies showing upregulation of gene expression in response to radiation in combination with QTP. Using mouse-specific primers for the corresponding murine genes we did not observe an upregulation of these genes in normal cells in the tumor microenvironment, thus indicating that this response was specific to GBM cells (**Fig. 2b**).

**Figure 2.**
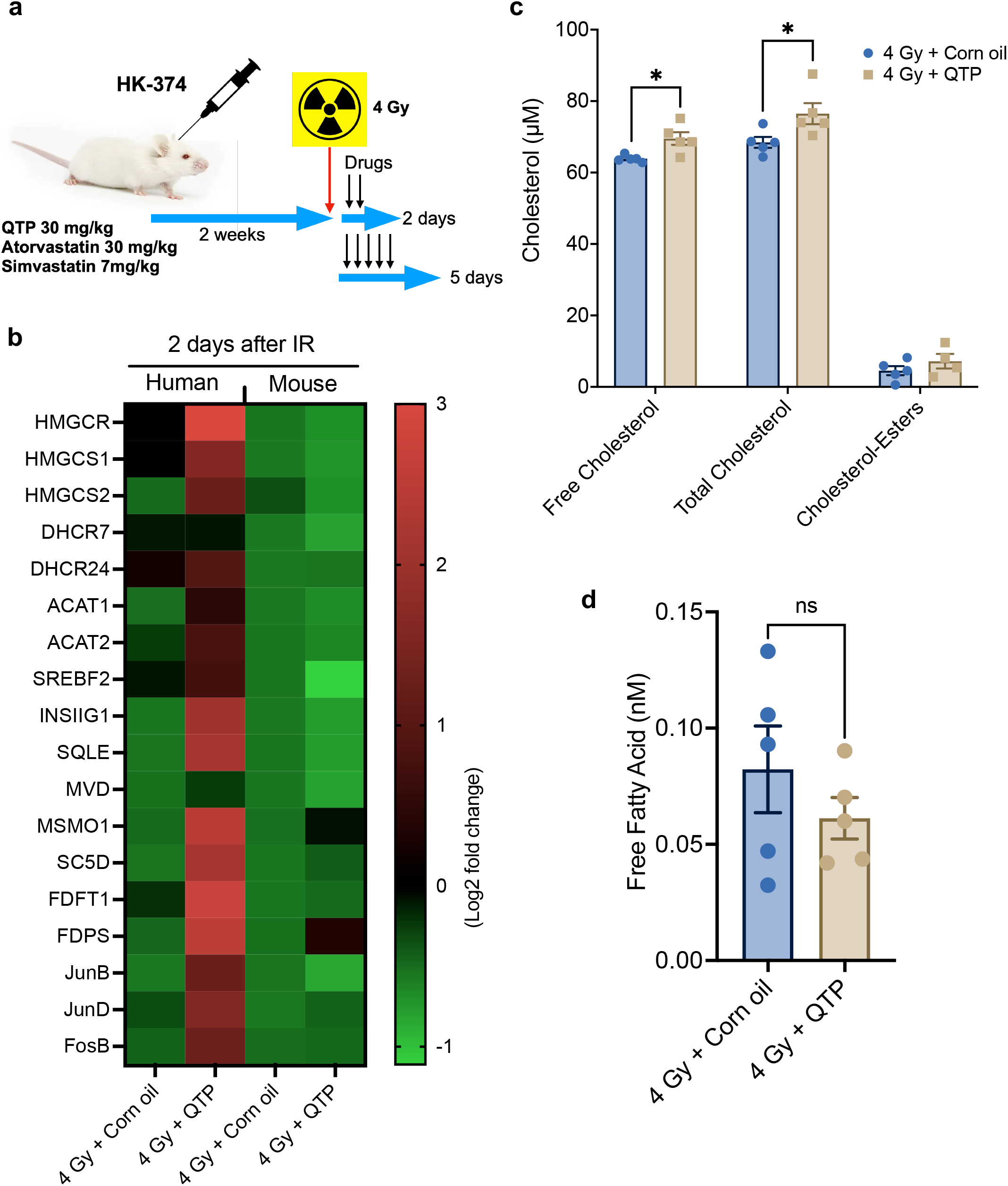
Upregulation of the mevalonate pathway in response to anti-cancer treatments *in vivo* is restricted to glioma cells. **(a)** Schematic of the experimental design underlying Figures 2 and 3. **(b)** Heatmap showing the results of quantitative RT-PCR for the cholesterol biosynthesis related genes in the tumor specimen harvested from the PDOX GBM mouse model at day 2 after the treatment of radiation (a single dose of 4 Gy) in combination with QTP (30 mg/kg) or solvent control using the human- and mouse-specific primers. **(c)** The levels of total, free cholesterol and cholesterol-esters in the tumors harvested from the PDOX GBM mouse model at day 2 after the treatments. **(d)** The concentration of free fatty acid in the tumors harvested from the PDOX GBM mouse model at day 2 after the treatments. All experiments have been performed with at least 3 biological independent repeats. *p-* values were calculated using Unpaired Student’s t-tests. * *p*-value < 0.05, ns: not significant.

In *in vitro* shotgun lipidomics experiments we previously demonstrated that radiation in combination with QTP upregulated cholesterol and free fatty acids production [3]. To test if these pathways were also functionally upregulated in response to the combination therapy *in vivo*, we next quantified cholesterol and free fatty acids levels in tumor tissues. Compared to radiation alone, the combination of radiation and QTP significantly upregulated free and total cholesterol levels in the tumor tissue (**Fig. 2c**) but did not affect free fatty acid levels (**Fig. 2d**), confirming that the mevalonate pathway was not only transcriptionally but also functionally activated. However, the increases in cholesterol synthesis were unexpectedly small, thus hinting that other branches of the mevalonate pathway could be involved in mediating the survival of GBM cells.

### Statins reduce treatment-induced upregulation of cholesterol biosynthesis in PDOXs

Statins are a class of FDA-approved drugs that target 3-hydroxy-3-methylglutaryl-CoA reductase (HMGCR), the rate-limiting enzyme in the mevalonate pathway. Designed to lower cholesterol levels in the periphery, data on blood-brain barrier (BBB) penetration in the literature is sparse and often conflicting. Our *in vivo* data indicated that the addition of statins to a combination of radiation and QTP could significantly improve median survival in mouse models of GBM [3]. Therefore, we next tested if statins could be detected in normal brain tissue and if statin treament would alter the mevalonate pathway in PDOXs.

We had previously used atorvastatin at a dose of 30 mg/kg in mice, and at this dose we were able to detect atorvastatin in brain tissue and plasma of mice with a brain to plasma ratio of 1.6 (**Fig. 3a**). *In silico* calculations predicted better BBB penetration for simvastatin **(Suppl. Fig. 2)** [11]. However, simvastatin is a prodrug with a short half-live and subject to first pass elimination [12, 13]. When we treated mice with simvastatin at 7 mg/kg, the murine equivalent dose [14] of the currently recommended human dose of 40 mg/day, mass spectrometry analysis of brain tissue and plasma could only detect plasma levels of simvastatin (**Fig. 3b**). However, both -atorvastatin and simvastatin-reduced free and total cholesterol but not free fatty acid levels in orthotopic gliomas (**Fig. 3c/d**) and prevented upregulation of gene expression of genes in the mevalonate pathway by radiation combined with QTP in tumor tissues (**Fig. 3e**). This indicated that both statins crossed the BBB in pharmacologically relevant concentrations or that there is sufficient diminution of the BBB in xenografts to let it it..

**Figure 3.**
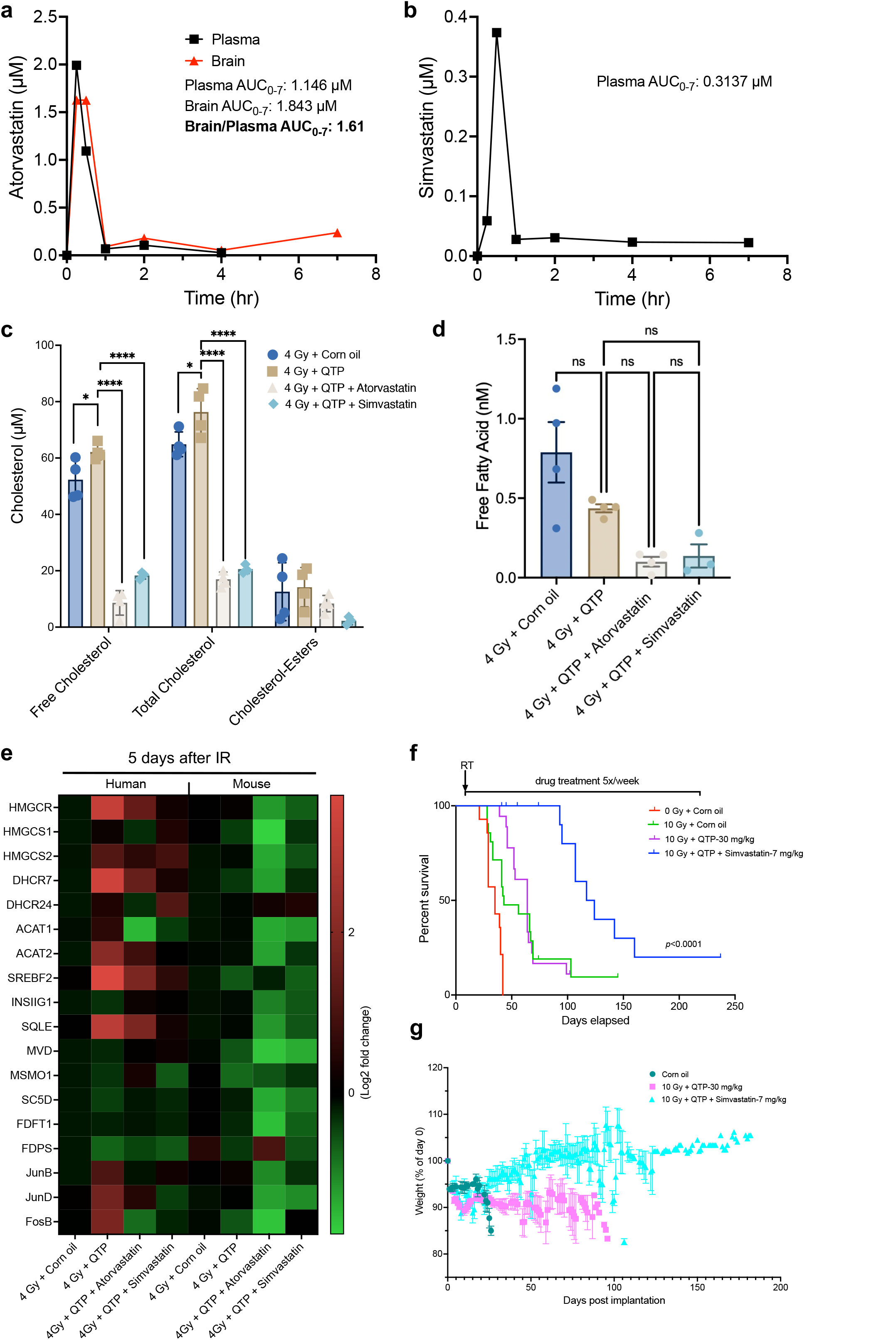
Statins reduce treatment-induced upregulation of cholesterol biosynthesis in PDOXs. **(a/b)** Brain and plasma levels of atorvastatin and simvastatin in C57BL/6 mice after a single injection (atorvastatin – 30 mg/kg, i.p.; simvastatin – 7 mg/kg, i.p.). **(c)** The levels of total, free cholesterol and cholesterol-esters in the tumors harvested from the PDOX GBM mouse model at day 5 after the treatment of radiation (a single dose of 4 Gy) in combination with QTP (30 mg/kg), atorvastatin (30 mg/kg) or simvastatin (7 mg/kg). **(d)** The concentration of free fatty acid in the tumors harvested from the PDOX GBM mouse model at day 5 after the treatments. **(e)** Heatmap showing the results of quantitative RT-PCR for the cholesterol biosynthesis related genes in the tumor specimen harvested from the PDOX GBM mouse model at day 5 after the treatments using the human- and mouse-specific primers. **(f)** Survival curves for NSG mice implanted intracranially with 3x10^5^ HK374-GFP-Luciferase glioma cells and grafted for 3 days. Mice were irradiated with a single fraction of 0 or 10 Gy and treated with corn oil or QTP (30 mg/kg, subQ, 5-day on/2-day off schedule) or triple combination of 10 Gy plus QTP and simvastatin (7 mg/kg, i.p., 5-day on/2-day off schedule) continuously until they reached the study endpoint. Log-rank (Mantel-Cox) test for comparison of Kaplan-Meier survival curves. **(g)** Weight curves for the NSG mice in different treatment groups. All experiments have been performed with at least 3 biological independent repeats. *p-*values were calculated using One-way ANOVA. * *p*-value < 0.05, **** *p*-value < 0.0001, ns: not significant.

### Statins improve median survival in mouse models of GBM undergoing fractionated irradiation

In previous *in vitro* studies we reported that the dopamine receptor antagonists trifluoperazine (TFP) and QTP prevented radiation-induced phenotype conversion of non-stem glioma cells into glioma stem cells, but induced gene expression of the mevalonate pathway in surviving cells, thereby creating a metabolic vulnerability that could be exploited with the use of atorvastatin to further improve median survival [3]. At 7 mg/kg, equivalent to the recommended human dose of 40 mg, simvastatin also significantly prolonged the median survival of glioma-bearing mice when combined with radiation and QTP (10 Gy + QTP: 64 days *vs.* 10 Gy + QTP + simvastatin: 120.5 days, *p*<0.0001, Log-rank test; **Fig. 3f**) and was well-tolerated with animals gaining weight during treatment (**Fig 3g**). We previously reported that the atypical dopamine receptor 2/3 antagonist and caseinolytic protease P (ClpP) activator ONC201 [15], now in clinical trials against pediatric glioma, also prevented radiation-induced phenotype conversion of non-stem glioma cells into glioma stem cells and prolonged median survival in mouse models of GBM [5]. Here we report that similar to TFP and QTP, ONC201 in combination with radiation also upregulates the mevalonate pathway, although with different kinetics, reflective of the longer half-life of ONC201 (**Fig. 4a**). Importantly, the addition of a statin significantly improved median survival compared to animals treated with radiation and ONC201 alone (10 Gy + ONC201 + atorvastatin *vs.* 10 Gy + ONC201, *p*=0.0193, Log-rank test; **Fig. 4b**), and led to smaller tumors or no signs of residual tumor in some animals as shown in H&E staining **(Fig. 4c)**.

**Figure 4.**
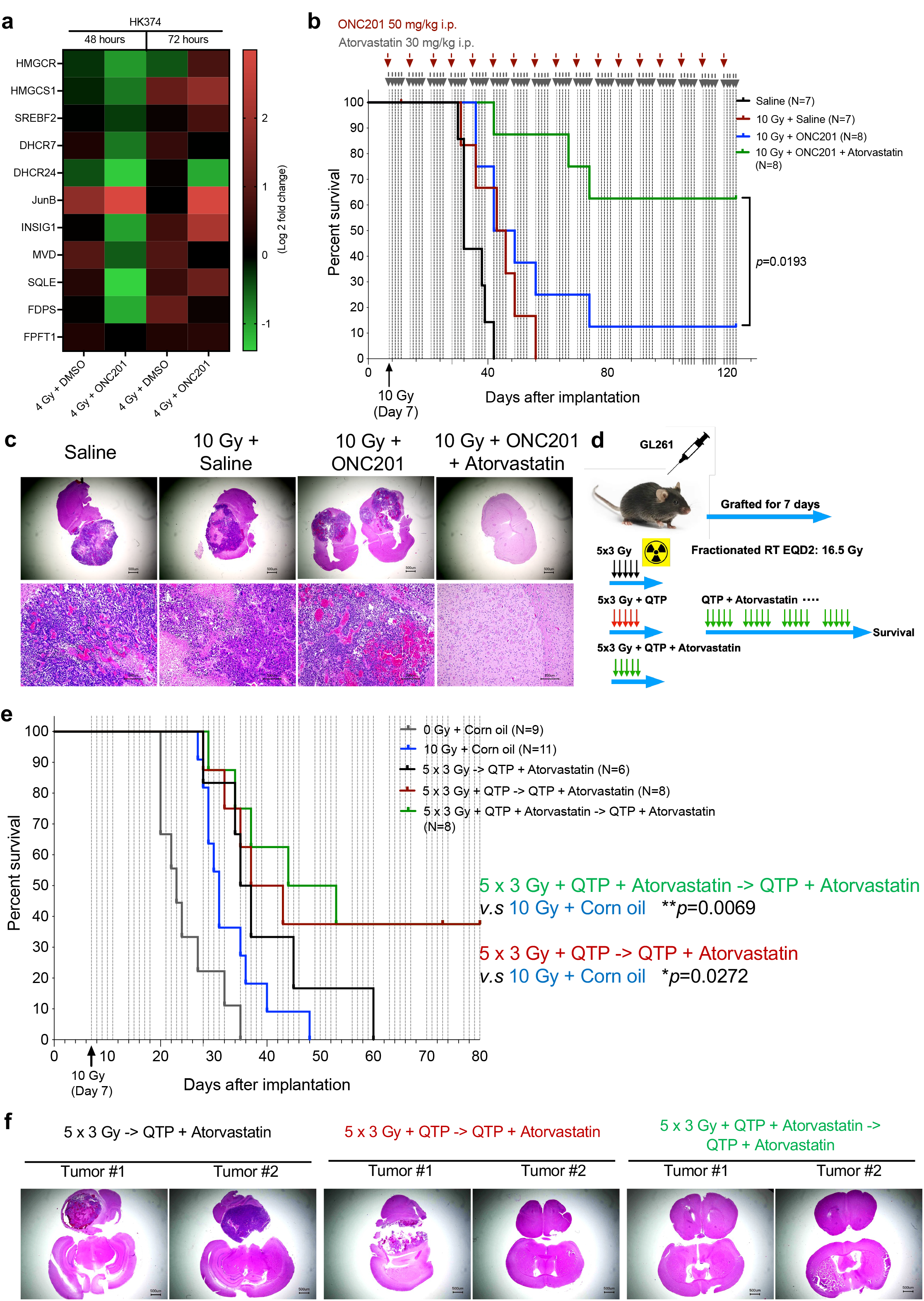
Statins improve median survival in mouse models of GBM undergoing fractionated irradiation. **(a)** Heat map showing the results of quantitative RT-PCR for the cholesterol biosynthesis related genes in HK374 cells treated with radiation (a single dose of 4 Gy) in the presence of absence of ONC201 (a single treatment of 2.5 µM) at 48 and 72 hours. **(b)** Survival curves for C57BL/6 mice implanted intracranially with 2x10^5^ GL261-GFP-Luciferase mouse glioma cells and grafted for 7 days. Mice were irradiated with a single fraction of 0 or 10 Gy and weekly treated with Saline or ONC201 (50 mg/kg, i.p.) or triple combination of 10 Gy plus ONC201 (weekly) and atorvastatin (30 mg/kg, i.p., 5-day on/2-day off schedule) continuously until they reached the study endpoint. Log-rank (Mantel-Cox) test for comparison of Kaplan-Meier survival curves. **(c)** H&E-stained coronal sections of the C57BL/6 mice brains implanted with GL261-GFP-Luc cells which were irradiated at 10 Gy and treated continuously with ONC201 in the presence of absence of atorvastatin until they met the criteria for study endpoint. **(d)** Schematic of the experimental design of fractionated irradiation in syngeneic mouse model of GBM. **(e)** Kaplan-Meier survival curves for C57BL/6 mice implanted intracranially with GL261-GFP-Luciferase mouse glioma cells and treated with either a single fraction of 0 or 10 Gy or 5 daily fractions of 3 Gy each and daily doses of either corn oil, QTP (30 mg/kg, subQ), or QTP plus atorvastatin (30 mg/kg, i.p.). After completion of the radiation treatment all animals were treated with QTP plus atorvastatin until they reached criteria for euthanasia. Log-rank (Mantel-Cox) test for comparison of Kaplan-Meier survival curves. **(f)** H&E stained coronal sections of the C57BL/6 mice brains from the groups of 5 x 3 Gy -> QTP + atorvastatin, 5 x 3 Gy + QTP -> QTP + atorvastatin and 5 x 3 Gy + QTP + atorvastatin -> QTP + atorvastatin.

Next, we repeated the experiment with fractionated irradiation (5 daily fractions of 3 Gy) in GL261 glioma bearing mice. In parallel, mice were treated with daily doses of either corn oil, QTP, or QTP plus atorvastatin. After completion of the radiation treatment all animals were treated with QTP plus atorvastatin until they reached criteria for euthanasia **(Fig. 4d)**. Kaplan-Meier estimates for the 3 treatment arms did not differ significantly but showed a trend for improved median survival for animals receiving QTP or QTP + atorvastatin during radiation treatment (**Fig. 4e**) in line with QTP preventing radiation-induced phenotype conversion of non-stem GBM cells into GSCs and atorvastatin affecting the mevalonate pathway in surviving cells. Histologically, the triple combination of radiation, QTP and atorvastatin led to smaller tumor sizes or no signs of residual tumor in some animals (**Fig. 4f**).

### Treatment-induced upregulation of the mevalonate pathway in glioma affects stemness through prenylation of Rac-1

The mevalonate pathway is the fundamental pathway in the cholesterol biosynthesis. While the uptake of exogenous cholesterol has been shown to be essential for glioma [16] other metabolites in the mevalonate pathway contribute to normal cell function through farnesylation and geranylgeranylation to the prenylation of small GTPases [17, 18]. To determine which of the pathway components’ upregulation contributes to glioma stemness, we employed an inhibitor-based approach (**Fig. 5a)**. Using clonal sphere- forming capacity assay, we found that inhibition of geranylgeranylation (**Fig. 5b**) but not inhibition of the cholesterol biosynthesis (**Fig. 5c/d**) or farnesylation (**Fig. 5e**) further reduced sphere formation when combined with radiation and QTP.

**Figure 5.**
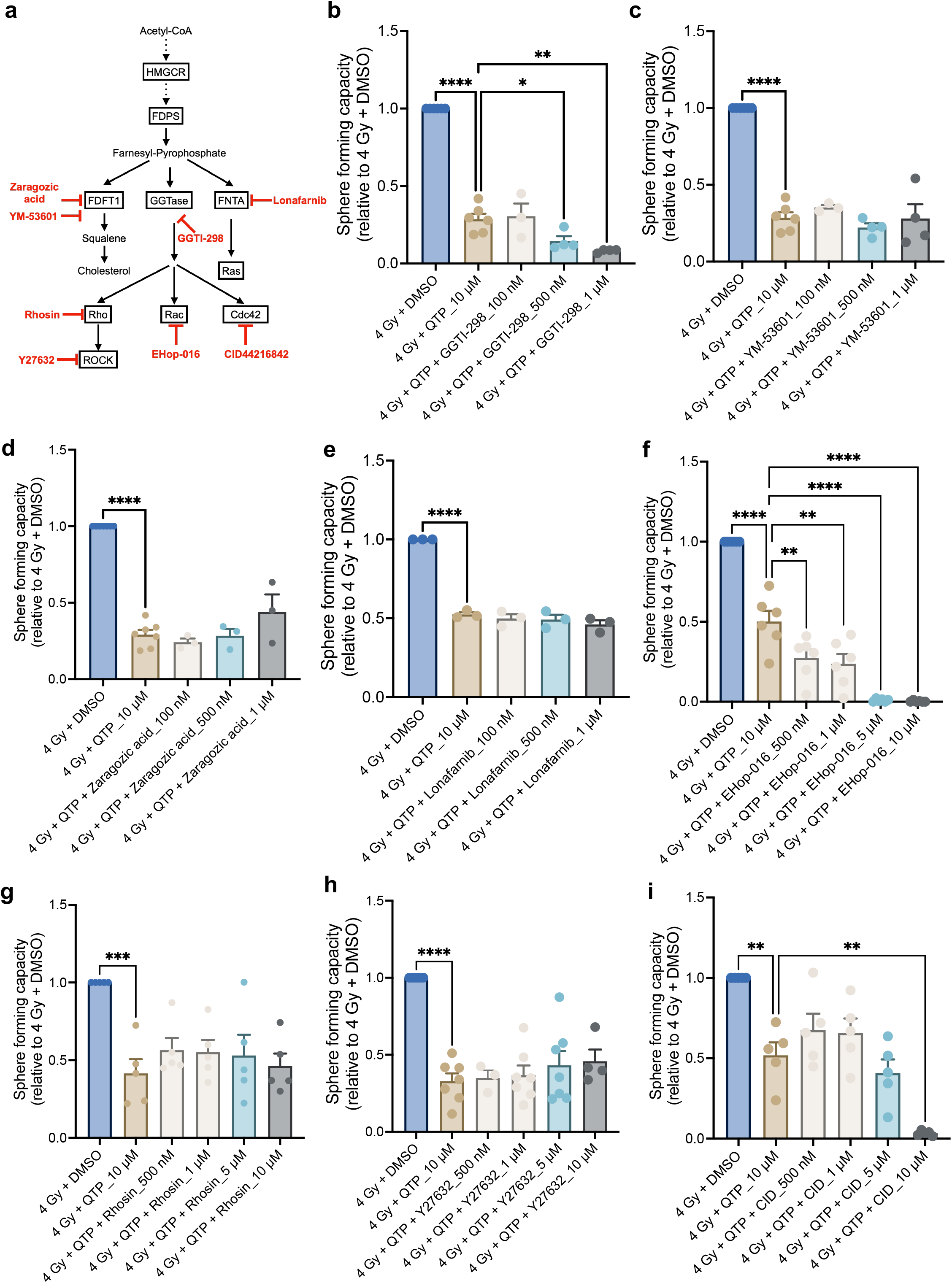
Identification of Rac1 as the potential contributor for maintaining the stemness of surviving glioma cells. (**a**) Cholesterol biosynthesis pathway and the key selected inhibitors. (**b-e**) Sphere-forming capacity of HK374 spheres treated either with GGTI-298 (GGTase inhibitor) or YM-53601 (squalene synthase inhibitor) or Zaragozic acid (squalene synthase inhibitor) or Lonafarnib (farnesyltransferase inhibitor) at 100, 500 nM, 1 µM concentrations when combined with radiation (a single dose of 4 Gy) and QTP (10 µM). (**f-i**) Sphere-forming capacity of HK374 spheres treated either with Ehop-016 (Rac GTPase inhibitor) or Rhosin (RhoA-specific inhibitor) or Y27632 (ROCK1/2 inhibitor) or CID44216842 (Cdc42-selective inhibitor) at 500 nM, 1, 5, 10 µM concentrations when combined with radiation and QTP. All experiments have been performed with at least 3 biological independent repeats. *p-*values were calculated using One-way ANOVA. * *p*-value < 0.05, ** *p*-value < 0.01, *** *p*-value < 0.001, **** *p*-value < 0.0001.

Since geranylgeranylation of Rho GTPases is fundamental for their downstream effects on cytoskeletal reorganization [19], which mediates the interaction of cancer stem cells (CSCs) with the tumor microenvironment, maintenance of stemness and CSC migration [20] we next tested which Rho GTPase would affect the self-renewal capacity of GBM cells (**Fig. 5a**). Inhibition of Rac GTPases (**Fig. 5f**) but not Rho or Cdc42 GTPases (**Fig. 5g-i**) significantly reduced sphere formation in a dose-dependent fashion when combined with radiation and QTP.

In agreement with our inhibitor studies, Rac-1 pulldown assays revealed a significant activation of Rac-1 after combined treatment with QTP and radiation (**Fig. 6a/b**). *In vitro*, surviving cells treated with radiation and QTP showing significantly increased migratory capacity (**Fig. 6c/d**), known to require remodeling of the cytoskeleton [21]. The addition of atorvastatin inhibited Rac-1 activation (**Fig. 6a**) and diminished the increased migratory capacity seen in cells surviving the combination treatment (**Fig. 6c/d**). Microtubules are part of the cytoskeleton, a structural network within the cell’s cytoplasm which helps to support cell shape, as well as cell migration and cell division. Using a microtubule stain we showed that the surviving cells treated with radiation and QTP had more filopodia (**Fig. 6e, white arrowheads)**, which support cell migration by promoting cell–matrix adhesiveness at the leading edge, as well as increased numbers of tunneling nanotubes (TNTs) (**Fig. 6e, yellow arrowheads).** TNTs are known to mediate protein and mitochondrial transfer between distant cells as part of the radiation response of GBM cells and are suspected to drive GBM growth and treatment resistance [22, 23]. This suggested more active Rac-1-mediated cytoskeleton remodeling in cells surviving the combination of radiation and QTP. Importantly, the observed effects on cytoskeleton remodeling were inhibited with the additional atorvastatin **(Fig. 6e)**.

**Figure 6.**
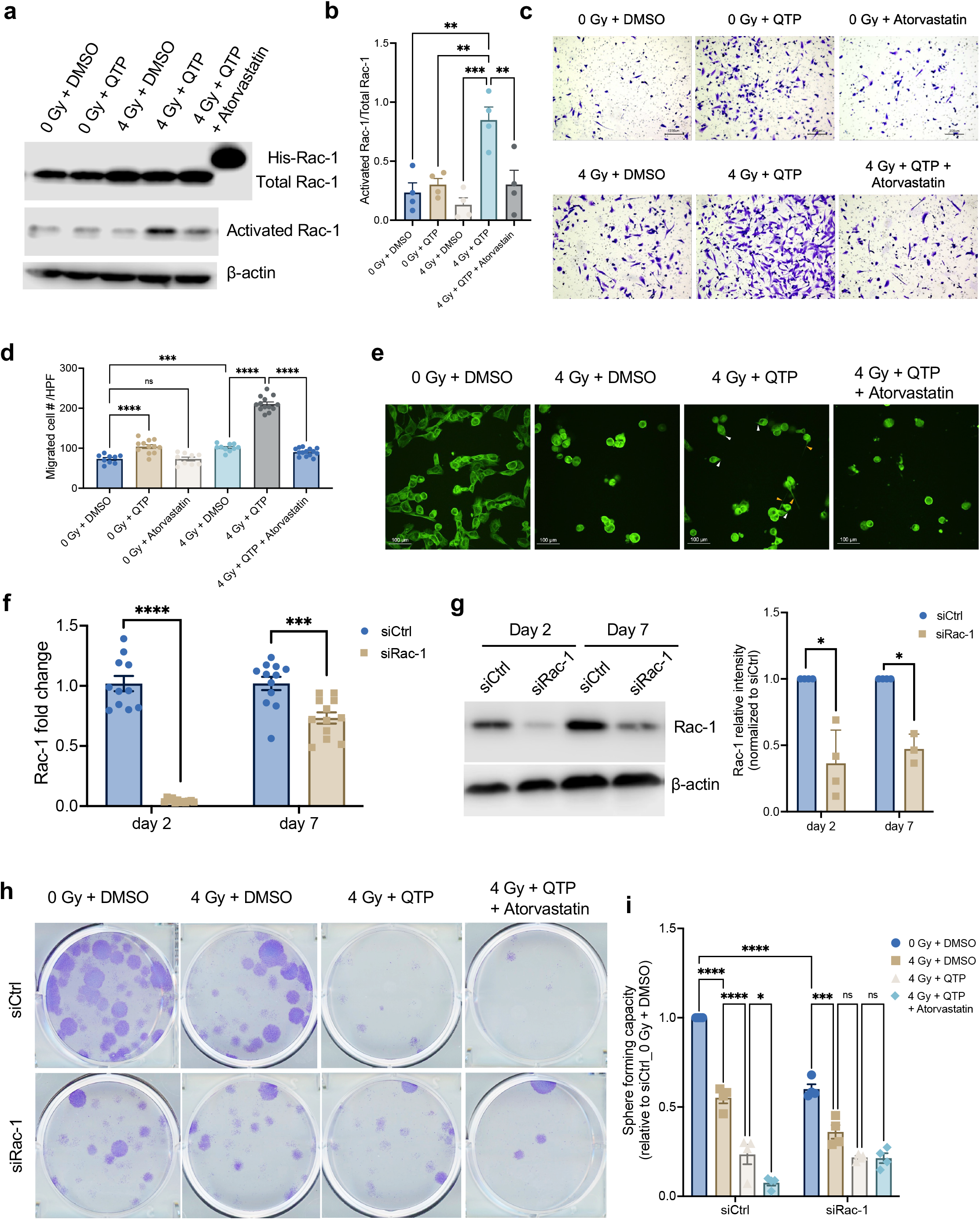
Treatment-induced upregulation of the mevalonate pathway in glioma affects stemness through prenylation of Rac1. **(a)** HK374 cells were treated with 0 or 4 Gy in the presence or absence of QTP (10 µM) and/or atorvastatin (1 µM) for 48 hours. The activated Rac1 was immunoprecipitated by 10 µg PAK-PBD agarose beads from the whole cell lysates and subjected to immunoblotting against Rac1, along with the total proteins. His-tagged Rac1 protein serves as the positive control. **(b)** The densitometry measurements of activated Rac1/total Rac1 using Image J. **(c)** Transwell migration assay of HK374 cells pre-treated with 0 or 4 Gy in the presence or absence of QTP (10 µM) and/or atorvastatin (1 µM) for 48 hours. **(d)** The quantification of migrated cells using Image J. **(e)** Confocal images of microtubules in HK374 cells treated with 0 or 4 Gy in the presence or absence of QTP (10 µM) and/or atorvastatin (1 µM) for 48 hours. White arrowheads: filopodia. Yellow arrowheads: tunneling nanotubes (TNTs). **(f/g)** Rac1 knock-down efficiency was evaluated at both mRNA (qRT-PCR) and protein (western blotting) levels at day 2 and day 7 after siRNA transfection. β-actin was used as the loading control and the densitometry measurements of Rac1 using Image J. **(h)** Clonogenic assay of siCtrl or siRac1 transfected HK374 cells treated with 0 or 4 Gy in the presence or absence of QTP (10 µM) and/or atorvastatin (1 µM) for 7 days. **(i)** Sphere-forming capacity of siCtrl or siRac1 HK374 spheres treated with 0 or 4 Gy in the presence or absence of QTP (10 µM) and/or atorvastatin (1 µM). All experiments have been performed with at least 3 biological independent repeats. *p-*values were calculated using One-way ANOVA for **b, d**; Unpaired Student’s t-tests for **f, g**; Two-way ANOVA for **i**. * *p*-value < 0.05, ** *p*-value < 0.01, *** *p*-value < 0.001, **** *p*-value < 0.0001, ns: not significant.

Lastly, we performed loss-of-function experiments to demonstrate that radiation/QTP-induced Rac-1 activation is indeed a driver for stemness maintenance in surviving GSCs. With the efficient transient knock down of Rac-1 in HK374 cells **(Fig. 6f/g)**, we tested its role in both, clonogenicity survival and self-renewal capacity of the cells surviving from radiation and/or QTP and the effect of atorvastatin. While treatment with radiation and QTP significantly reduced the formation of adherent clonal colonies (**Fig. 6h**), sphere-forming capacity (**Fig. 6i)** and stem cell frequency **(Tables 1 and 2)**, additional treatment with atorvastatin did not further diminish these three parameters after knock-down of Rac-1, thus suggesting that all three traits are Rac-1-driven in cells surviving radiation and QTP.

**Table 1.** Confidence intervals for stem cell frequency (%).

| **Groups** | **Lower** | **Estimate** | **Upper** |
| --- | --- | --- | --- |
| siCtrl_0 Gy + DMSO | 7.246377 | 9.90099 | 13.45895 |
| siCtrl_4 Gy + DMSO | 3.584229 | 4.901961 | 6.697924 |
| siCtrl_4 Gy + QTP | 0.785546 | 1.079914 | 1.483239 |
| siCtrl_4 Gy + QTP + Atorvastatin | 0.403877 | 0.555247 | 0.762893 |
| siRac-1_0 Gy + DMSO | 4.608295 | 6.329114 | 8.613264 |
| siRac-1_4 Gy + DMSO | 2.083333 | 2.857143 | 3.913894 |
| siRac-1_4 Gy + QTP | 1.48368 | 2.040816 | 2.800336 |
| siRac-1_4 Gy + QTP + Atorvastatin | 0.816327 | 1.122334 | 1.541545 |

**Table 2.** Pairwise tests for differences in stem cell frequencies.

| **Group 1** | **Group 2** | **Chisq** | **DF** | **Pr(>Chisq)** |
| --- | --- | --- | --- | --- |
| siCtrl_0 Gy + DMSO | siCtrl_4 Gy + DMSO | 8.81 | 1 | 0.003 |
| siCtrl_4 Gy + DMSO | siCtrl_4 Gy + QTP | 41 | 1 | 1.51e-10 |
| siCtrl_4 Gy + QTP | siCtrl_4 Gy + QTP + Atorvastatin | 7.86 | 1 | 0.00507 |
| siCtrl_0 Gy + DMSO | siRac-1_0 Gy + DMSO | 3.64 | 1 | 0.0562 |
| siCtrl_4 Gy + DMSO | siRac-1_4 Gy + DMSO | 4.95 | 1 | 0.0261 |
| siRac-1_0 Gy + DMSO | siRac-1_4 Gy + DMSO | 10.7 | 1 | 0.00105 |
| siRac-1_4 Gy + DMSO | siRac-1_4 Gy + QTP | 1.92 | 1 | 0.165 |
| siRac-1_4 Gy + QTP | siRac-1_4 Gy + QTP + Atorvastatin | 6.06 | 1 | 0.0138 |

## Discussion

To date, surgery and radiotherapy remain the most effective treatment modalities against GBM. Temozolomide, the only chemotherapeutic agent added to the standard-of-care during the past two decades [24, 25], only marginally improves patient outcome and the median and long-term survival rates of patients suffering from GBM remain unacceptably low. Attempts to increase the tumoricidal efficacy of radiotherapy through e.g., alternative fractionation schemes or radiosensitizers have largely failed [26], in part due to the dispersion of glioma cells into the normal parenchyma beyond the visible tumor and because many compounds lacked BBB penetration or increased normal tissue toxicity. Likewise, targeted therapies had little effects as GBMs escape through utilization of alternate signaling pathways [27].

The DNA-damaging effects of radiation unfold in milliseconds after irradiation and the repair of DNA double-strand breaks is completed within the first few hours. Compounds that increase the amount of initial DNA damage or interfere with its repair mainly operate during this short timeframe. While it is desirable to eliminate all malignant cells this way, current therapies against GBM fail in achieving this goal and almost all patients succumb to progressive or recurrent disease. Over the past two decades it became evident that a small population of cancer stem cells, relatively resistant to radio- and chemo-therapy, maintains the growth of tumors [28–30] and shows enrichment after treatment [2, 31, 32]. Recurrent or progressing gliomas are likely caused by GSCs that survive these sublethal treatments and the pathways that drive them to proliferate, invade, and repopulate the tumor are not necessarily identical with the pathways that helped them to survive the genotoxic insults.

With the discovery of radiation-induced phenotype conversion of non-stem cancer cells into cancer stem cells we have reported a novel major factor for radiation treatment failure [33]. In an unbiased phenotypic high-throughput screen we identified compounds that -unlike TMZ, which enhances it-prevented this unwanted effect of radiation, and we demonstrated that repurposing FDA-approved drugs identified in this screen prolonged median survival in PDOX models of glioma [3, 4].

The mevalonate pathway is one of the key metabolic pathways known to be dysregulated in GBM [34]. Statins, developed to lower cholesterol levels in the periphery target HMGCR, the rate-limiting enzyme in this pathway and affect glioma biology in multiple ways, from interfering with the dependence of GBM on cholesterol to inhibition of farnesylation of Ras or geranylgeranylation of GTPases with effects on cell proliferation and survival, cell cycle progression, migration and invasion [35]. Recognized as a potential target, the mevalonate pathway has been probed in several preclinical and clinical studies repurposing statins against brain tumors with largely disappointing results. While statins showed robust anti-tumor activities *in vitro* [35–37], previous rodent pharmacokinetic studies were not able to achieve therapeutic drug concentrations in the central nervous system (CNS) [38, 39]. A recent clinical phase II study, which added atorvastatin to the current standard-of-care, radiotherapy and TMZ, showed no beneficial effects on progression-free or overall survival when compared to historic controls treated with radiotherapy and TMZ [40]

Clinically, the biodistribution of TMZ into the brain is less than 20% and peak concentrations after single doses of 75-200 mg/m^2^ were predicted to reach only 1.8-3.7 µg/ml or 9.2 to 19.1 µM [41]. At these concentrations TMZ has only minimal if any effects *in vitro* and most preclinical studies use TMZ in the high micromolar to millimolar range [42, 43]. Although TMZ is a known activator of the MAPK cascade, our *in vitro* data did not indicate effects of TMZ on the mevalonate pathway unless drug concentrations were raised to 1 mM. The lack of upregulation of the mevalonate pathway in response to low TMZ concentrations agreed with the lack of clinical benefits of statins in combination with radiation and TMZ, which was not surprising given that 23% of the human body’s cholesterol are located in the brain [44] and that GBM cells rely on exogenous cholesterol and not on cholesterol biosynthesis [16].

In contrary, dopamine receptor antagonists activate the MAPK cascade via GSK-3 at low concentrations and synergize with radiation by targeting tumor cell plasticity and glioma stem cells [4, 5]. Designed as psychotropic drugs they easily cross the blood-brain barrier and reach therapeutic concentrations in the CNS. Here we show that dopamine receptor antagonists robustly created a metabolic vulnerability in surviving cells through upregulation of the mevalonate pathway *in vivo*.

Tumors outpace cell loss with proliferation, thereby creating a need for macromolecular building blocks to not only replicate DNA but also to double the cellular mass before and during mitosis. To serve this need, cancer cells utilize aerobic glycolysis, known as the Warburg effect, to channel glucose into the biosynthesis of macromolecules including the mevalonate pathway to produce cholesterol. The cell cycle and the mevalonate pathway are tightly intervened with non-sterol isoprenoids initiating the transition from G_1_- to S-phase and cholesterol being required during the G_2_- and M phase of the cell cycle. Very high cholesterol concentration are toxic for cells and cholesterol biosynthesis is limited by a negative feedback loop [45]. And since GBM cells primarily rely on uptake of exogenous cholesterol [16] it is not surprising that the radiation/QTP-induced upregulation of the mevalonate pathway reported here only slightly increased cholesterol biosynthesis in GBM cells.

Consequently, the inhibition of this branch of the mevalonate pathway did not affect the self-renewal of surviving GBM cells beyond the effects of radiation and dopamine receptor antagonists. Instead of upregulating cholesterol production, surviving cells increased their migratory and invasive capacity through activation of Rac-1, a major regulator of cytoskeleton reorganization. A role for Rac-1 in stemness and tumorigenicity had been previously reported for U87MG and U373 glioma cells [46] and our results confirmed these findings with a significant reduction in clonogenic survival, sphere-forming capacity and glioma stem cell frequency when Rac-1 was knocked down. Likewise, the treatment of cells with a geranylation inhibitor or inhibition of Rac-1 synergized with radiation and QTP in reducing the sphere-forming capacity of the cells.

We conclude that the upregulation of the mevalonate pathway is part of the immediate-early response to glioma treatments that converge in activating the MAPK cascade, effectively diminish the GSC pool and interfere with glioma cell plasticity. Surviving GSCs rely on activation of Rac-1 through the mevalonate pathway to maintain stemness and to repopulate the tumors. This generates a metabolic vulnerability that can be exploited through the addition of statins to further improve the efficacy of radiotherapy. Furthermore, upregulation of the mevalonate pathway seemed to be restricted to tumor tissue, indicating the presence of a therapeutic window for statins in combination with radiation and repurposed, FDA-approved dopamine receptor antagonists.

These effects on the mevalonate pathway are not seen after treatment with radiation and TMZ, the current standard of care, and the addition of statins is not expected to provide an anti-tumor effect in this situation.

## Methods

### Cell lines

Primary human glioma cell lines were established at UCLA as described in [29]; Characteristics of specific gliomasphere lines can be found in [47]. Primary GBM cells were propagated as gliomaspheres in serum-free conditions in ultra-low adhesion plates in DMEM/F12, supplemented with SM1 Neuronal Supplement (#05177, STEMCELL Technology, Kent, WA) EGF (#78006, STEMCELL Technology), bFGF (#78003, STEMCELL Technology) and heparin (1,000 USP Units/mL, NDC0409-2720-31, Lake Forest, IL) as described previously [29, 47, 48]. GL261 cells were cultured in log-growth phase in DMEM supplemented with 10% fetal bovine serum, penicillin and streptomycin (P/S). All cells were grown in a humidified atmosphere at 37°C with 5% CO_2_. The unique identity of all patient-derived specimens was confirmed by DNA fingerprinting (Laragen, Culver City, CA). All lines were routinely tested for mycoplasma infection (#G238, Applied biological Materials, Ferndale, WA).

### Animals

6–8-week-old C57BL/6 mice, or NOD-*scid* IL2Rgamma^null^ (NSG) originally obtained from The Jackson Laboratories (Bar Harbor, ME) were re-derived, bred and maintained in a pathogen-free environment in the American Association of Laboratory Animal Care-accredited Animal Facilities of Department of Radiation Oncology, University of California, Los Angeles, in accordance with all local and national guidelines for the care of animals. 2x10^5^ GL261-GFP-Luc or 3x10^5^ HK374-GFP-Luc cells were implanted into the right striatum of the brains of mice using a stereotactic frame (Kopf Instruments, Tujunga, CA) and a nano-injector pump (Stoelting, Wood Dale, IL). Injection coordinates were 0.5 mm anterior and 2.25 mm lateral to the bregma, at a depth of 3.0 mm from the surface of the brain. Weight of the animals was recorded daily. Tumors were grown for 3 days for HK374 cells and 7 days for GL261 cells with successful grafting confirmed by bioluminescence imaging. Mice that developed neurological deficits or lost 20% of their body weights requiring euthanasia were sacrificed.

### Drug treatment

For *in vivo* studies, after confirming tumor grafting via bioluminescence imaging, mice implanted with HK374 cells were injected subcutaneously with quetiapine (QTP; #KS-1099, Key Organics, Cornwall, UK) at 30 mg/kg and/or intraperitoneally with simvastatin at 7 mg/kg on a 5-days on / 2-days off schedule until they reached euthanasia endpoints. QTP was dissolved in acidified PBS (0.4% Glacial acetic acid) at a concentration of 5 mg/ml. Simvastatin (#S1792, Selleck Chemicals, Houston, TX) was dissolved in corn oil containing 2.5% DMSO at a concentration of 5 mg/ml. Mice bearing GL261 tumors were injected intraperitoneally on a weekly basis either with ONC201 at 50 mg/kg and/or intraperitoneally with atorvastatin (#10493, Cayman Chemical, Ann Arbor, MI) at 30 mg/kg until they reached the euthanasia endpoint. ONC201 was kindly provided by Oncoceutics, Inc. (Philadelphia, PA) and dissolved in sterile saline at a concentration of 5.5 mg/ml. Atorvastatin was dissolved in corn oil containing 2.5% DMSO at a concentration of 5 mg/ml.

To determine the optimal timing of the drug treatments during fractionated radiotherapy. Mice bearing GL261 tumors received 5 daily fractions of 3 Gy. In parallel, mice were treated with daily doses of either saline, QTP (30 mg/kg, subQ), or QTP + atorvastatin (30 mg/kg, i.p.). After completion of the radiation treatment all animals were treated with QTP + atorvastatin until they reached criteria for euthanasia.

To explore the activation of MAPK pathway *in vitro*, HK374 cells were serum starved and the following day treated with a single dose of 10 Gy in the absence or presence of QTP (10 µM), temozolomide (1 mM; #14163, Cayman Chemical) or Vincristine (250 nM; #HY-N0488, MedChem Express, Monmouth Junction, NJ). 2 hours after the irradiation, cell lysates were harvested for western botting.

To determine, which part of the mevalonate pathways synergizes with radiation and QTP to affect self-renewal, HK374 cells were plated under sphere forming conditions in an *in-vitro* limiting dilution assay. Cells were irradiated with a single dose of 4 Gy in the presence of QTP (10 µM) and treated either with zaragozic acid (squalene synthase inhibitor; #17452, Cayman Chemical,), GGTI-298 (GGTase inhibitor; #16176, Cayman Chemical), YM-53601 (squalene synthase inhibitor; #18113, Cayman Chemical), Lonafarnib (farnesyltransferase inhibitor, #SML1457, Sigma, St. Louis, MO) at 100 nM, 500 nM, and 1 µM or Rhosin (RhoA-specific inhibitor; #555460, Sigma), Y27632 (ROCK1/2 inhibitor; #S1049, Selleck Chemicals), EHop-016 (Rac GTPase inhibitor; #S7319, Selleck Chemicals), CID44216842 (Cdc42-selective inhibitor; #S6000, Selleck Chemicals) at 500 nM, 1 µM, 5 µM and 10 µM. All drugs were dissolved in DMSO at 10 mM for stock and were replenished daily for 5 days.

### Migration Assay

HK374 cells were treated with 10 µM QTP, 1 µM atorvastatin, or QTP and atorvastatin and irradiated with 0 or 4 Gy. 48 hours after irradiation, cells were serum-starved for 12 hours. 5x10^5^ cells were then plated into 12-well Transwell^®^ insert with 8 µm pore size (Corning, New York, NY). After 24 hours, membranes were washed with PBS, cells fixed with 10% formalin, stained with 1% crystal violet, and counted using ImageJ.

### *In-vitro* limiting dilution assay

For the assessment of self-renewal capacity *in vitro*, HK374 cells were seeded at clonal densities under serum-free conditions into non-tissue-culture-treated 96-well plates in DMEM/F12 media, supplemented with SM1 Neuronal Supplement, EGF, bFGF and heparin. The next day, the cells were treated with drugs and one hour later irradiated with a single dose of 4 Gy. Growth factors were supplemented every two days. Glioma spheres were counted 10 days later and presented as the percentage of the initial number of cells plated.

### Irradiation

Cells were irradiated with at room temperature using an experimental X-ray irradiator (Gulmay Medical Inc. Atlanta, GA) at a dose rate of 5.519 Gy/min. Control samples were sham-irradiated. The X-ray beam was operated at 300 kV and hardened using a 4 mm Be, a 3 mm Al, and a 1.5 mm Cu filter and calibrated using NIST-traceable dosimetry. Corresponding controls were sham irradiated.

For *in vivo* irradiation experiments, mice were anesthetized prior to irradiation with an intra-peritoneal injection of 30 µL of a ketamine (100 mg/mL, Phoenix, MO) and xylazine (20 mg/mL, AnaSed, Lake Forest, IL) mixture (4:1) and placed on their sides into an irradiation jig that allows for irradiation of the midbrain while shielding the esophagus, eyes, and the rest of the body. For survival experiments, animals implanted with GL261 cells received a single dose of 10 Gy on day 7 or 5 fractions of 3 Gy each starting 7 days after tumor implantation. Animals injected with HK374 glioma specimen received a single dose of 10 Gy on day 3 after tumor implantation.

### Quantitative Reverse Transcription-PCR

Total RNA was isolated using TRIZOL Reagent (Invitrogen, Waltham, MA) cDNA synthesis was carried out using the SuperScript Reverse Transcriptase IV (Invitrogen). Quantitative PCR was performed in the QuantStudio^TM^ 3 Real-Time PCR System (Applied Biosystems, Carlsbad, CA, USA) using the PowerUp^TM^ SYBR^TM^ Green Master Mix (Applied Biosystems). *C*_t_ for each gene was determined after normalization to PPIA and ΔΔ*C*_t_ was calculated relative to the designated reference sample. Gene expression values were then set equal to 2^−ΔΔCt^ as described by the manufacturer of the kit (Applied Biosystems). All PCR primers were synthesized by Invitrogen with PPIA as the housekeeping gene (for primer sequences see **Supplementary Table 2**).

### Cholesterol and free fatty acid quantification

3x105 HK374-GFP-Luc cells were intracranially implanted and grafted for 2 weeks to achieve the appropriate size of tumor to start with. Mice were irradiated with a single dose of 4 Gy and treated with daily with either corn oil, QTP (30 mg/kg), QTP + atorvastatin (30 mg/kg) or QTP + simvastatin (7 mg/kg). After 2 and 5 days of drug treatments, the mice were euthanized, and brain tumor samples were collected and weighted. PBS (20 µl/mg) was added to mince the brain tumor tissues with a pellet pestle® tissue grinder (#749521-1590, DWK Life Sciences, Rockwood, TN). Equal volumes of homogenized tumor specimens were subsequently subjected to Cholesterol/Cholesterol Ester-GLo^TM^ assay (#J3190, Promega, Madison, WI) and Free Fatty Acid assay (ab65341, Abcam, Cambridge, UK) following the manufacturers’ instructions.

### Western Blotting

HK374 and HK217 cells were serum starved overnight and the following day treated with 10 μM QTP, 1 mM TMZ, or 250 nM Vincristine for one hour and then irradiated with a single dose of 10 Gy. Two hours after irradiation, the cells were lysed in 150 μl of ice-cold RIPA lysis buffer (10 mM Tris-HCl (pH 8.0), 1 mM EDTA, 1 % Triton X-100, 0.1 % Sodium Deoxycholate, 0.1 % SDS, 140 mM NaCl, 1 mM PMSF) containing protease inhibitor (#A32955, Thermo Fisher Scientific, Waltham, MA) and phosphatase inhibitor (#A32957, Thermo Fisher Scientific). The protein concentration in each sample was determined by BCA protein assay (Thermo Fisher Scientific) and samples were denaturated in 4X Laemmli sample buffer (Bio-Rad) containing 10% β-mercaptoethanol for 10 mins at 95°C. Equal amounts of protein were loaded onto 10% SDS-PAGE gels (1X Stacking buffer - 1.0 M Tris-HCl, 0.1% SDS, pH 6.8, 1X Separating buffer - 1.5 M Tris-HCl, 0.4% SDS, pH 8.8) and were subjected to electrophoresis in 1X Running buffer (12.5 mM Tris-base, 100 mM Glycine, 0.05% SDS), initially at 50 V for 30 min followed by 100 V for 2 hours. Samples were then transferred onto 0.45 μm nitrocellulose membrane (Bio-Rad, Hercules, CA) for 2 hours at 80 V. Membranes were blocked in 1X TBST (20 mM Tris-base, 150 mM NaCl, 0.2% Tween-20) containing 5% bovine serum albumin (BSA) for 30 min and then incubated with primary antibodies against *p*-ERK (#4370S, 1:1000, Cell Signaling, Danvers, MA), *p*-P38 (#4511S, 1:1000, Cell Signaling), t-ERK (#5013S, 1:1000, Cell Signaling), t-P38 (#8690S, 1:1000, Cell Signaling), Rac1 (#ARC03, 1:500, Cytoskeleton, Denver, CO), β-actin (#3700S, 1:1000, Cell Signaling) in 1X TBST containing 5% BSA overnight at 4°C with gentle rocking. Membranes were then washed three times for 5 min each with 1X TBST and incubated with secondary antibodies, 1:2000 anti-mouse or anti-rabbit horseradish peroxidase (HRP; Cell Signaling) in TBST for 2 hours at room temperature with gentle rocking. Membranes were washed again three times for 5 min each with 1X TBST. Pierce ECL Plus Western Blotting Substrate (Thermo Fisher Scientific) was added to each membrane and incubated at room temperature for 5 min. The blots were then scanned with Odyssey Fc Imaging system (LI-COR, Lincoln, NB). β-actin was used as a loading control. The ratio of the gene of interest over its endogenous control was calculated and expressed as relative intensity.

### Rac1 pulldown assay

HK374 cells were serum starved overnight, treated with QTP (10 µM) or QTP + atorvastatin (1 µM) and irradiated with a single dose of 4 Gy one hour after the drug treatment. 48 hours after the treatment, the cells were lysed in 250 µl cell lysis buffer supplemented with protease inhibitors and subjected to a pull-down assay for activated Rac1 (#BK035, Cytoskeleton). Briefly, the protein concentration in each sample was determined using the BCA protein assay (Therma Fisher Scientific) and 500 µg cell lysate was incubated with PAK-PBD beads at 4°C on a rotator for one hour. The beads were pelleted by centrifugation, washed with 500 µl wash buffer for two times, resuspended in 20 μl of 2x Laemmli sample buffer and boiled at 100°C for 5 minutes. The pull-down protein samples were then subjected to western blotting with the whole cell lysate and His-tagged Rac1 protein (#RC01, Cytoskeleton) serving as the controls.

### siRNA and Extreme Limiting Dilution Analysis (ELDA)

HK374 cells were seeded into 6-well tissue-culture plates and transfected with scramble siRNA or Rac1 siRNA by incubation with RNAiMAX-siRNA Lipofectamine duplex (#13778075, Thermo Fisher Scientific) in Opti-MEM medium overnight at 37°C with 5% CO_2._ The next day, cell culture medium was changed back to regular DMEM + 10% FBS

+ 1% P/S. Two days after the transfection, the cells were trypsinized and plated into the non-tissue-culture-treated 96-well plates in DMEM/F12 media, supplemented with SM1 Neuronal Supplement, EGF, bFGF and heparin. The next day, the cells were treated with DMSO or QTP (10 µM) or QTP + atorvastatin (1 µM) and irradiated with a single dose of 4 Gy one hour after the drug treatment. Growth factors were supplemented every two days. Glioma spheres were counted 5 days later (day 7 after siRNA transfection) and presented as the percentage to the initial number of cells plated. The glioma stem cell frequency was calculated using the ELDA software [49]. Both protein and RNA samples were collected at day 2 and day 7 after transfection. Knockdown efficiency was confirmed by RT-PCR and western blotting.

### Clonogenic assay

Two days after the siRNA transfection, HK374 cells were trypsinized and plated at a density of 100 cells per well in a 6-well plate. The next day, the cells were treated with DMSO or QTP (10 µM) or QTP + atorvastatin (1 µM) and irradiated with a single dose of 4 Gy one hour after the drug treatment. The colonies were fixed and stained with 0.1% crystal violet 5 days later (day 7 after siRNA transfection). Colonies containing at least 50 cells were counted in each group.

### Microtubule stain

HK374 cells were plated into 35 mm dish (No. 1.5 Coverslip, 10 mm Glass Diameter, Poly-D-Lysine coated; #P35GC-1.5-10-C, MatTek Ashland, MA) and the following day treated with QTP (10 µM) or QTP + atorvastatin (1 µM) and irradiated with a single dose of 4 Gy one hour after the drug treatment. 48 hours later, cells were stained with ViaFluor® 488 live cell microtubule stains (#70062, Biotium, Fremont, CA) following manufacturer’s instructions. Specifically, the cells were washed with PBS and incubated with medium containing probes (0.5 x ViaFluor® dye + 100 µM verapamil) at 37°C for 30 minutes. Fluorescent images were then acquired using a confocal microscope (Nikon A1, Melville, NY).

### Mass spectrometry

Sample Preparation: Whole blood from mice was centrifuged to isolate plasma. Atorvastatin or simvastatin was isolated by liquid-liquid extraction from plasma: 50 µL plasma was added to 2 µL internal standard and 100 µL acetonitrile. Mouse brain tissue was washed with 2 mL cold saline and homogenized using a tissue homogenizer with fresh 2 mL cold saline. Atorvastatin or simvastatin was then isolated and reconstituted in a similar manner by liquid-liquid extraction: 100 µL brain homogenate was added to 2 µL internal standard and 200 µL acetonitrile. The samples were centrifuged, supernatant removed and evaporated by a rotary evaporator and reconstituted in 100 µL 50:50 water:acetonitrile.

Atorvastatin or Simvastatin Detection: Chromatographic separations were performed on a 100 x 2.1 mm Phenomenex Kinetex C18 column (Phenomenex, Torrence, CA) using the 1290 Infinity LC system (Agilent, Santa Clara, CA). The mobile phase was composed of solvent A: 0.1% formic acid in Milli-Q water, and B: 0.1% formic acid in acetonitrile. Analytes were eluted with a gradient of 5% B (0-4 min), 5-99% B (4-32 min), 99% B (32-36 min), and then returned to 5% B for 12 min to re-equilibrate between injections. Injections of 20 µL into the chromatographic system were used with a solvent flow rate of 0.10 mL/min.

Mass spectrometry was performed on a 6460 triple quadrupole LC/MS system (Agilent). Ionization was achieved by using positive ion electrospray ionization and data acquisition was made in multiple reaction monitoring mode. Atorvastatin was monitored with the transition from m/*z* 559.2→250 with fragmentor settings at 45 V and a collision energy of 17 and an accelerator voltage of 4V, and for Simvastatin m/*z* 419.2→198.2 with fragmentor settings at 85 V and a collision energy of 5 and an accelerator voltage of 4 V. Atorvastatin brain concentrations were adjusted by 1.4% of the mouse brain weight for the residual blood in the brain vasculature as described by Dai *et al*. [50].

### Statistics

Unless stated otherwise all data shown are represented as mean ± standard error mean (SEM) of at least 3 biologically independent experiments. A *p*-value of ≤0.05 in an unpaired two-sided *t*-test, One-way or Two-way ANOVA indicated a statistically significant difference. For Kaplan-Meier estimates a *p*-value of 0.05 in a log-rank test indicated a statistically significant difference.

## Availability of data and materials

All data and methods are included in the manuscript. Patient-derived cell lines will be made available upon reasonable request to the corresponding author.

---

**Supplementary Figure 1.**
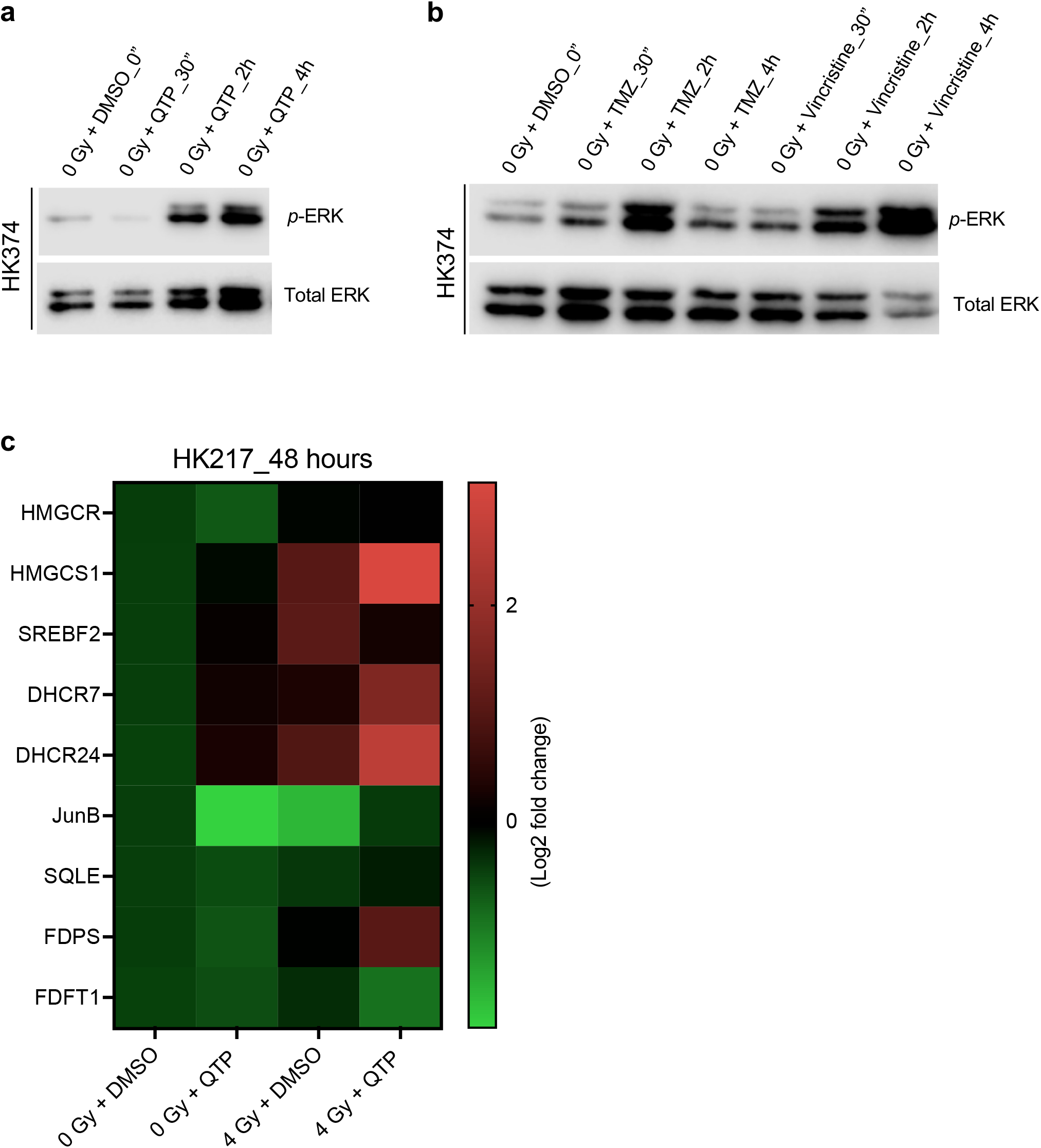

**Supplementary Figure 2.**
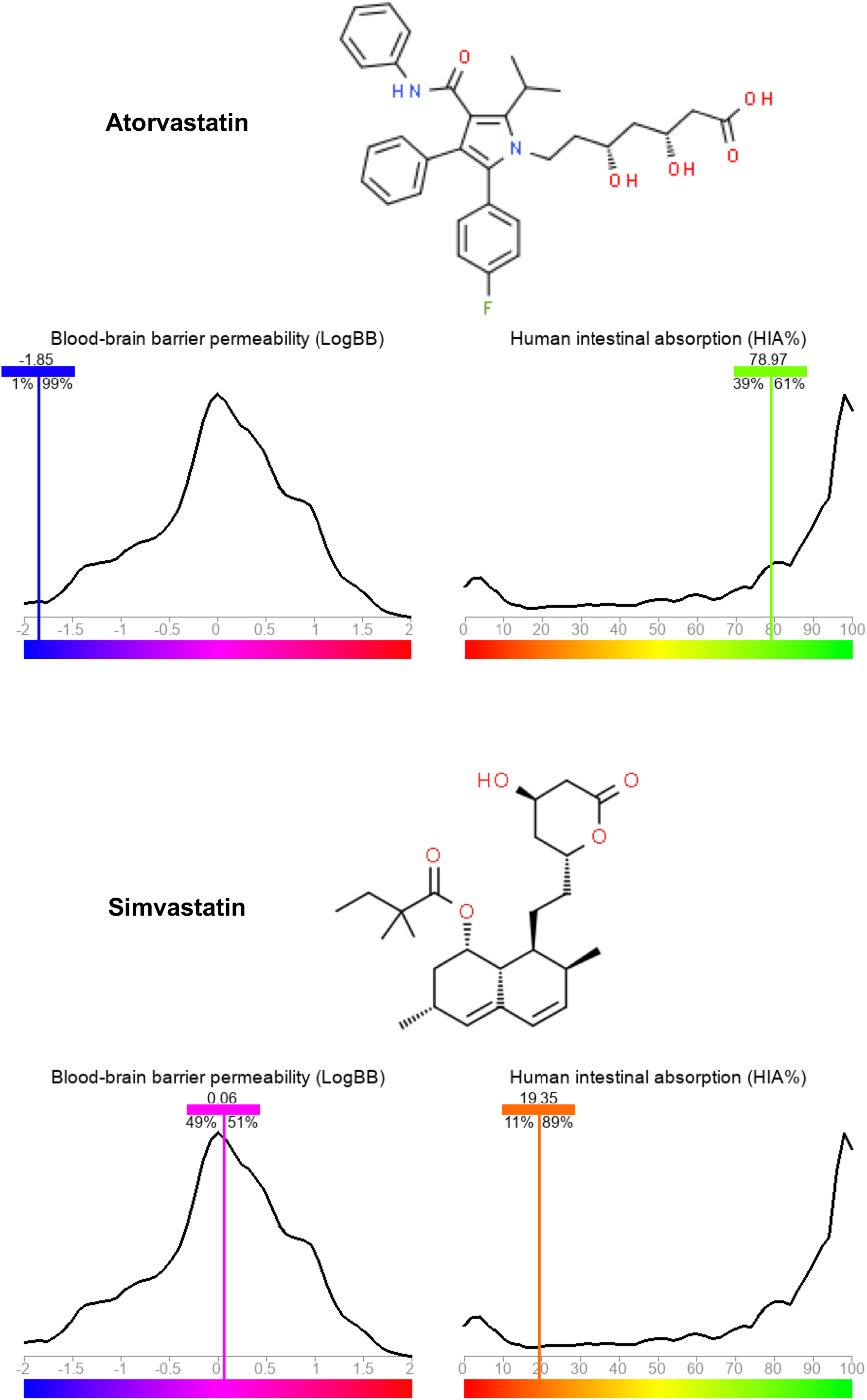

