## Supplemental Material for "Activation of the mevalonate pathway in response to anti-cancer treatments drives glioblastoma recurrences through activation of Rac-1"

#### Table of contents:


- Supplementary Figures
- Supplementary Tables

#### Supplementary Figures

##### Supplementary Figure 1.

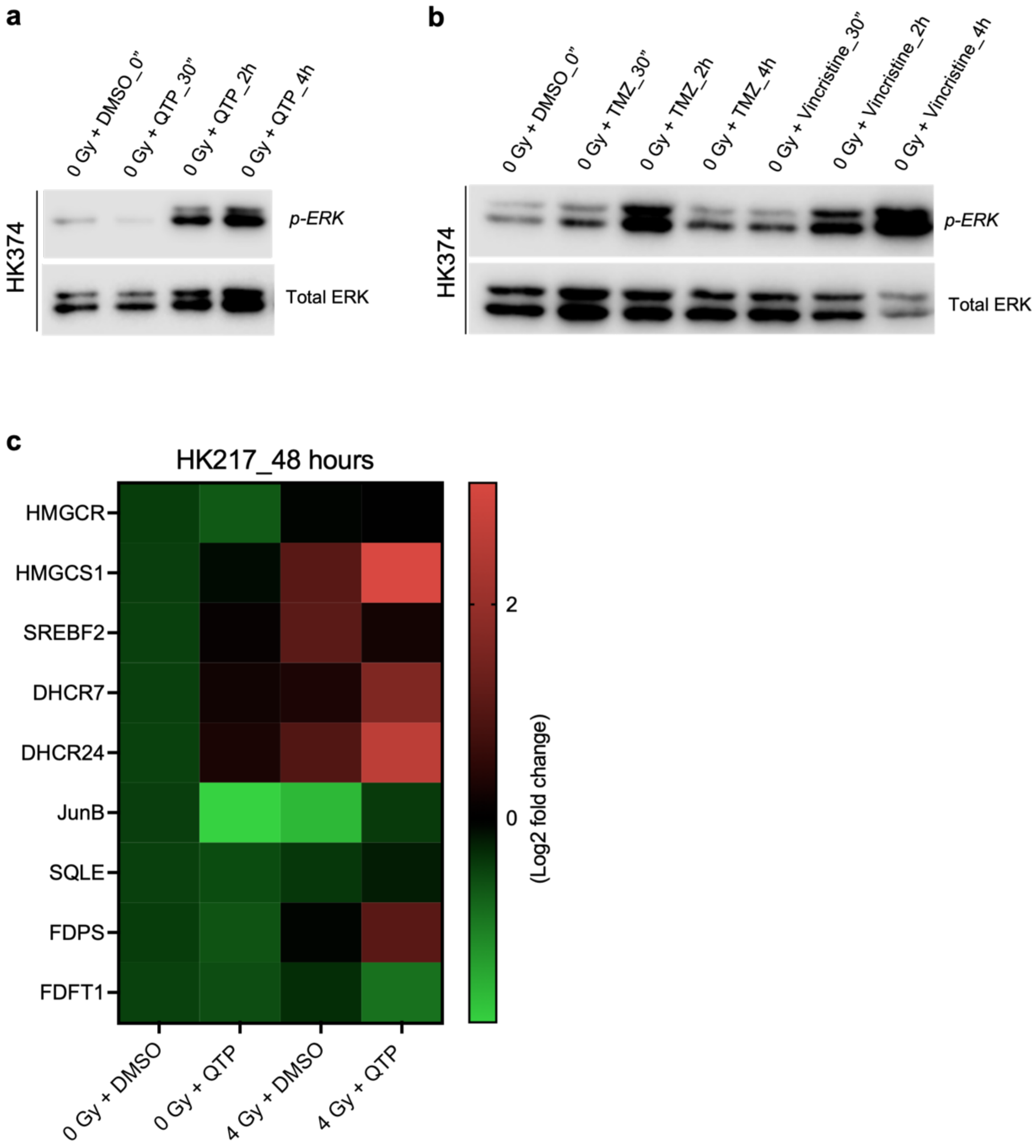


**Supplementary Figure 1.** **(a/b)** Western blotting of *p*-ERK and total ERK in HK374 cells treated with QTP (10 μM), TMZ (1 mM) or Vincristine (250 nM) for 30 minutes, 2 hours, and 4 hours, with the solvent DMSO-treated cells as the control. **(c)** Heatmap showing the results of quantitative RT-PCR for the cholesterol biosynthesis related genes in HK217 cells treated with radiation (a single dose of 4 Gy) in the presence of absence of QTP (10 μM) for two consecutive days. All experiments have been performed with at least 3 biological independent repeats.

#### Supplementary Figure 2

**Atorvastatin**

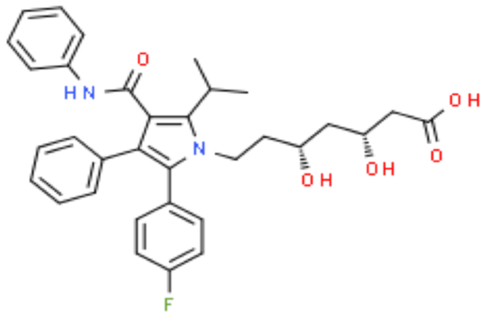


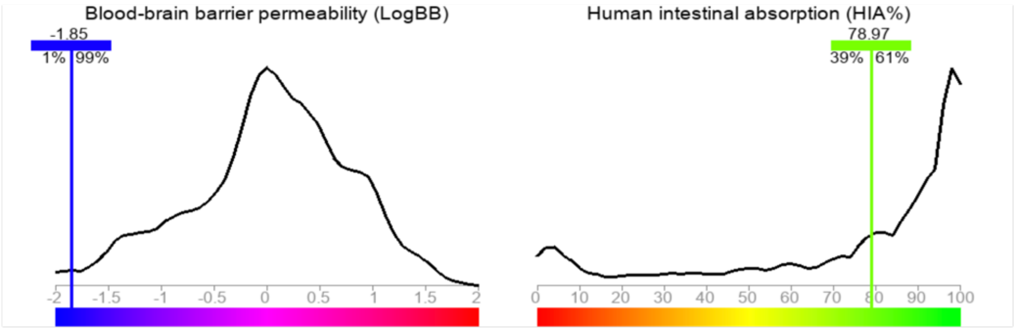


**Simvastatin**

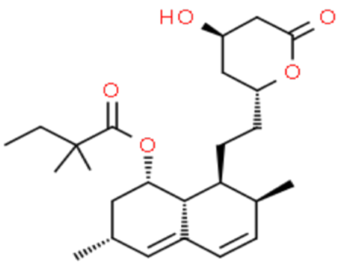


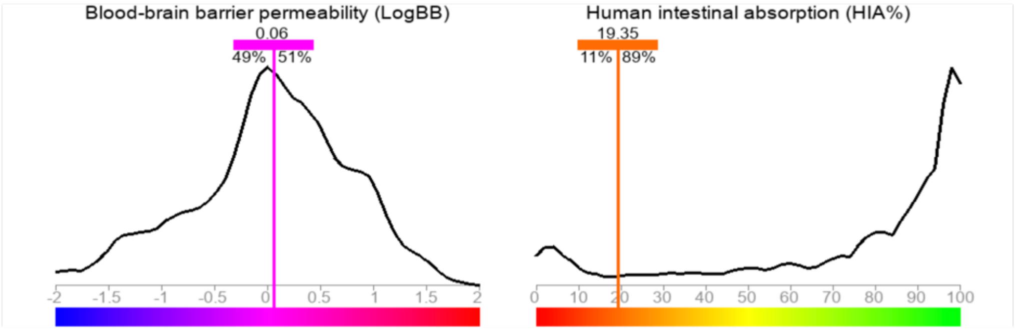


**Supplementary Figure 2.** Structures of atorvastatin and simvastatin and the predicted blood-brain barrier permeability (LogBB) and intestinal absorption (HIA%) by the integrated web service for predicting ADMET parameters of drugs, which is being developed in the Laboratory of Medicinal Chemistry, Department of Chemistry, Lomonosov Moscow State University.

### Supplementary Tables

#### Supplementary Table 1

##### Patient demographics and TCGA-classification of GBM subtypes.

| Cell line | Origin | Age | Gender | TCGA subtype | Culture P53 CN | EGFRvIII | PTEN | MGMT |
|---|---|---|---|---|---|---|---|---|
| HK374 | Primary GBM | 45 | Male | classical | Loss "mosaic" | Positive | Positive | not methylated |
| HK217 | Primary GBM | 81 | Male | proneural | wt | Negative | Negative | methylated |

**Supplementary Table 2**

| Species | Gene name | Primer sequence (5’–3’) |
|---|---|---|
| **Human** | HMGCR | Forward: TGATTGACCTTTCCAGAGCAAG<br>Reverse: CTAAAATTGCCATTCCACGAGC |
| **Human** | HMGCS1 | Forward: CATTAGACCGCTGCTATTCTGTC<br>Reverse: TTCAGCAACATCCGAGCTAGA |
| **Human** | HMGCS2 | Forward: TCCCTTTACCTCTCCACTCAC<br>Reverse: CCATAAGAGAAGGCACCAATCC |
| **Human** | DHCR7 | Forward: GCTGCAAAATCGCAACCCAA<br>Reverse: AGCTTTGGTTCTCTGCCTG |
| **Human** | DHCR24 | Forward: GCCGCTCTCGCTTATCTTCG<br>Reverse: GTCTTGCTACCCTGCTCCTT |
| **Human** | ACAT1 | Forward: CGGGCTAACTGATGTCTACAA<br>Reverse: CAAATTTCCCAGCTTCCCATG |
| **Human** | ACAT2 | Forward: CCCAGAACAGGACAGAGAATG<br>Reverse: AGCTTGGACATGGCTTCTATG |
| **Human** | SREBF2 | Forward: AACGGTCATTCACCCAGGTC<br>Reverse: GGCTGAAGAATAGGAGTTGCC |
| **Human** | INSIG1 | Forward: CCTGGCATCATCGCCTGTT<br>Reverse: AGAGTGACATTCCTCTGGATCTG |
| **Human** | SQLE | Forward: GATGATGCAGCTATTTTCGAGGC |

| | | |
|---|---|---|
| | | Reverse: CCTGAGCAAGGATATTCACGACA |
| **Human** | MVD | Forward: CGTGGCATCGGTGAACAACT<br>Reverse: GTGTAGGCTAGGCAGGCATA |
| **Human** | MSMO1 | Forward: TATGCTGGTTCTCGGCATCAT<br>Reverse: CCAAAAATTCGATCCCACCATGT |
| **Human** | SC5D | Forward: CATACGTGTATCCAGCCAC<br>Reverse: AAGAACAGTGCAACAGTAAGA |
| **Human** | FDFT1 | Forward: CCACCCCGAAGAGTTCTACAA<br>Reverse: TGCGACTGGTCTGATTGAGATA |
| **Human** | FDPS | Forward: TGTGACCGGCAAAATTGGC<br>Reverse: GCCCGTTGCAGACACTGAA |
| **Human** | JunB | Forward: GGACACGCCTTCTGAACG<br>Reverse: CGGAGTCCAGTGTGGTTTG |
| **Human** | JunD | Forward: CCTCAGCCACGTCAACAG<br>Reverse: CACCCTCTCCAAGTCCG |
| **Human** | FosB | Forward: AGCTAAATGCAGGAACCGG<br>Reverse: ACCAGCACAAACTCCAGAC |
| **Human** | Rac1 | Forward: GGTGAATCTGGGCTTATGGG<br>Reverse: TCAGGATACCACTTTGCACG |
| **Human** | PPIA | Forward: ATGCTGGACCCAACACAAAT<br>Reverse: TCTTTCACTTTGCCAAACACC |
| **Mouse** | HMGCR | Forward: GCCCTCAGTTCAAATTCACAG<br>Reverse: TTCCACAAGAGCGTCAAGAG |

| | | |
|---|---|---|
| **Mouse** | HMGCS1 | Forward: TGTTCTCTTACGGTTCTGGC<br>Reverse: AAGTTCTCGAGTCAAGCCTTG |
| **Mouse** | HMGCS2 | Forward: GTACCTTGAACGAGTGGATGAG<br>Reverse: GGTGGGATTTTAAGCAGATGC |
| **Mouse** | DHCR7 | Forward: TTATTCCTGGCTTCCTGACTTC<br>Reverse: CAGAGGATGTGGGTAATGAGC |
| **Mouse** | DHCR24 | Forward: AGAACTACCTGAAGACAAACCG<br>Reverse: GAAGAGGTAGCGGAAGATGG |
| **Mouse** | ACAT1 | Forward: AGCACACTGAACGATGGAG<br>Reverse: CGCAAGTGGAAAATCAATGGG |
| **Mouse** | ACAT2 | Forward: CTGGAGGCATGGAGAATATGAG<br>Reverse: CATGTGGTAGTTGTGAAAGGC |
| **Mouse** | SREBF2 | Forward: CCCTATTCCATTGACTCTGAGC<br>Reverse: CACATAAGAGGATTCGAGAGCG |
| **Mouse** | INSIG1 | Forward: GATTACCATCGCCTTCCTAGC<br>Reverse: CGTCCTATGTTTCCCACTGTG |
| **Mouse** | SQLE | Forward: CCCCAAAACACAAAATCCTCAG<br>Reverse: GCAATGCCAAGAAAAGTCCAC |
| **Mouse** | MVD | Forward: GCTCCGAATCCTTATCCTTGTG<br>Reverse: GGGTCATCTCCTTCATGCG |
| **Mouse** | MSMO1 | Forward: ATTTCCTGCACAGACTCCTTC<br>Reverse: AGAATCAGGGTTTCCAAGGG |
| **Mouse** | SC5D | Forward: CCGTCTCACTGTTCCTGC |

| | | |
|---|---|---|
| | | Reverse: GCCCCTATGAATCCAGTAGATC |
| **Mouse** | FDFT1 | Forward: GTGTGGGATGGCAGAATTTG<br>Reverse: GGCAGAGAATAGACGAGAAAGG |
| **Mouse** | FDPS | Forward: TCTTTCTACCTGCCTATTGCG<br>Reverse: CTCCAAAGAGATCAAGGTAGTCG |
| **Mouse** | JunB | Forward: GGACACGCCTTCTGAGAG<br>Reverse: GAGTCCAGTGTGTGAGCTG |
| **Mouse** | JunD | Forward: AACAGAAAGTCCTCAGCCAC<br>Reverse: AGTCTCGAAAGAGTCCGGG |
| **Mouse** | FosB | Forward: AGTCTCAGTACCTGTCTTCGG<br>Reverse: CACGAGCCACTGAAGATCC |
| **Mouse** | GAPDH | Forward: AGGTCGGTGTGAACGGATTTG<br>Reverse: CCCTGGCACATGAATCCTGG |
